# Emergence of distributed working memory in a human brain network model

**DOI:** 10.1101/2023.01.26.525779

**Authors:** Mengli Feng, Abhirup Bandyopadhyay, Jorge F. Mejias

**Affiliations:** Cognitive and Systems Neuroscience Group, Swammerdam Institute for Life Sciences, University of Amsterdam, 1090GE Amsterdam, the Netherlands

## Abstract

Working memory is a fundamental cognitive function which allows to transiently store and manipulate relevant information in memory. While it has been traditionally linked to activity in specific prefrontal cortical areas, recent electrophysiological and imaging evidence has shown co-occurrent activities in different brain regions during working memory. To dissect the mechanisms behind the emergence of such distributed working memory activity in the human brain, we built and analyzed a detailed, data-constrained model of the human brain combining multiple sources of large-scale brain data. The model reveals the emergence of distributed working memory patterns which rely on long-range synaptic projections, differing from classical models where working memory emerges due to local recurrent connectivity. When compared to existing evidence regarding areas linked to working memory, our model explains almost 80% of the observed variability across brain areas. Our results predict important differences between the distributed working memory in human and non-human primates, and propose a potential solution for the debate regarding the role of early visual areas in working memory encoding. This work provides a novel approach to simulate the human brain which combines the richness of large-scale brain dynamics with the insight of mechanistic models of cognitive functions.

## Introduction

Working memory is a flexible cognitive function which allows our brains to temporarily store and manipulate sensory information, and it is essential for cognitive tasks involving complex functional behavior such as decision making, reasoning, and learning. Working memory has traditionally been associated with elevated activity in frontal and prefrontal areas of the brain: electrophysiological explorations in non-human primates (NHPs) have specifically suggested an important role of dorsolateral prefrontal cortex (dlFPC)^1–6^, with positron emission tomography (PET) and functional magnetic resonance imaging (fMRI) studies in humans also highlighting the role of prefrontal activity for different types of working memory (such as visual or verbal tasks) in the human brain^7–10^. In more recent years, the study of working memory has been reinvigorated by the exploration of novel neurophysiological mechanisms to transiently sustain information in cortical networks, including the consideration of calcium-dependent synaptic contributions^11–13^ linked to activity-silent mechanisms^14,15^, time-varying neural representations^16–18^ and oscillatory processes^19,20^.

Both traditional electrophysiological studies in NHPs and computational models have continued to put the focus of working memory on prefrontal areas such as dlPFC. However, evidence from human neuroimaging studies and systematic multi-region recordings in NHPs^21–24^ and even in recent large-scale electrophysiological recordings in rodents^25^, have crystalized the idea that activity linked to working memory is found in multiple brain regions, often co-activated. For example, fMRI studies have shown that working memory tasks also activate other brain regions, such as the parietal cortex^26^, the medial temporal lobe^27^ or the cingulate cortex^28^. Additionally, studies have shown that different working memory tasks activate different brain regions, suggesting that there is not a single "working memory area" in the brain^29–31^. Despite all available evidence, particularly for the human brain, computational works have only recently proposed potential mechanisms for the emergence of distributed working memory for the case of NHPs^32^ and rodents^33^; the question remains unexplored for the human brain.

In this work, we took on the challenge of building a data-constrained large-scale computational model of the human brain to study the emergence of distributed working memory patterns in humans. To build such a model, we combined three complementary large-scale datasets: a structural human connectome comprising 180 left-hemisphere neocortical areas or regions-of-interest (ROIs) defined as in previous work^34,35^, an MRI-derived contrast ratio of T1-to-T2 weighted (T1w:T2w) maps which serves as an estimation of the hierarchical relationships between the 180 areas^35,36^, and a detailed brain-wide mapping of NMDA receptor density obtained from PET imaging^37^. This data allowed us to constrain the model not only at the level of long-range projections between brain areas, but also at the local level to regulate the strength of synaptic couplings in an area-specific way. The model was able to display the emergence of self-sustained activity triggered by visual input and distributed across multiple brain regions. In contrast to classical models of local attractors, such elevated activity originated primarily from long-range brain interactions (since local synaptic strength was too weak to produce multistability on its own^32^), and presented a significant level of robustness to distractors. Our model was able to explain up to 78% of the working memory-related activity reported across a wide range of experimental literature, and predicted substantial differences with respect to existing models of distributed working memory in NHPs. Finally, the model provided a plausible solution to the controversy of the involvement of early visual areas in working memory maintenance. Overall, our work denotes a first step in the mechanistic study of cognitive functions like working memory in human brain network models.

## Results

To build a detailed, data-constrained computational model of the human brain and explore the generation of distributed working memory, we combined (i) human connectivity from multimodal neuroimaging data from the Human Connectome Project (HCP)^34,35^, (ii) the intracortical myelin content for each area, measured from MRI-derived contrast ratio of T1-to-T2 weighted (T1w:T2w) maps and used to estimate the hierarchical relationship between areas^35,36^, and (iii) a PET-derived brain map of NMDA receptor density^37^. Using these three complementary datasets, we constrained our model at different levels (for details, see Methods). Briefly, the model consisted of a network of 180 neocortical areas (or ROIs, see Table S1) distributed across all lobes of the left hemisphere of the human brain (Fig. 1A). Areas were connected via inter-areal projections whose strength was constrained by the human connectivity data from previous work^34,35^ (Fig. S1A). The overall connectivity strength across the entire network was modulated by a parameter in the model, the global coupling strength (G). Each area was modelled using a simplified circuit model^38^ with two excitatory populations, selective to the two possible visual stimuli considered (labelled as A and B respectively), and one inhibitory population.

**Figure 1:**
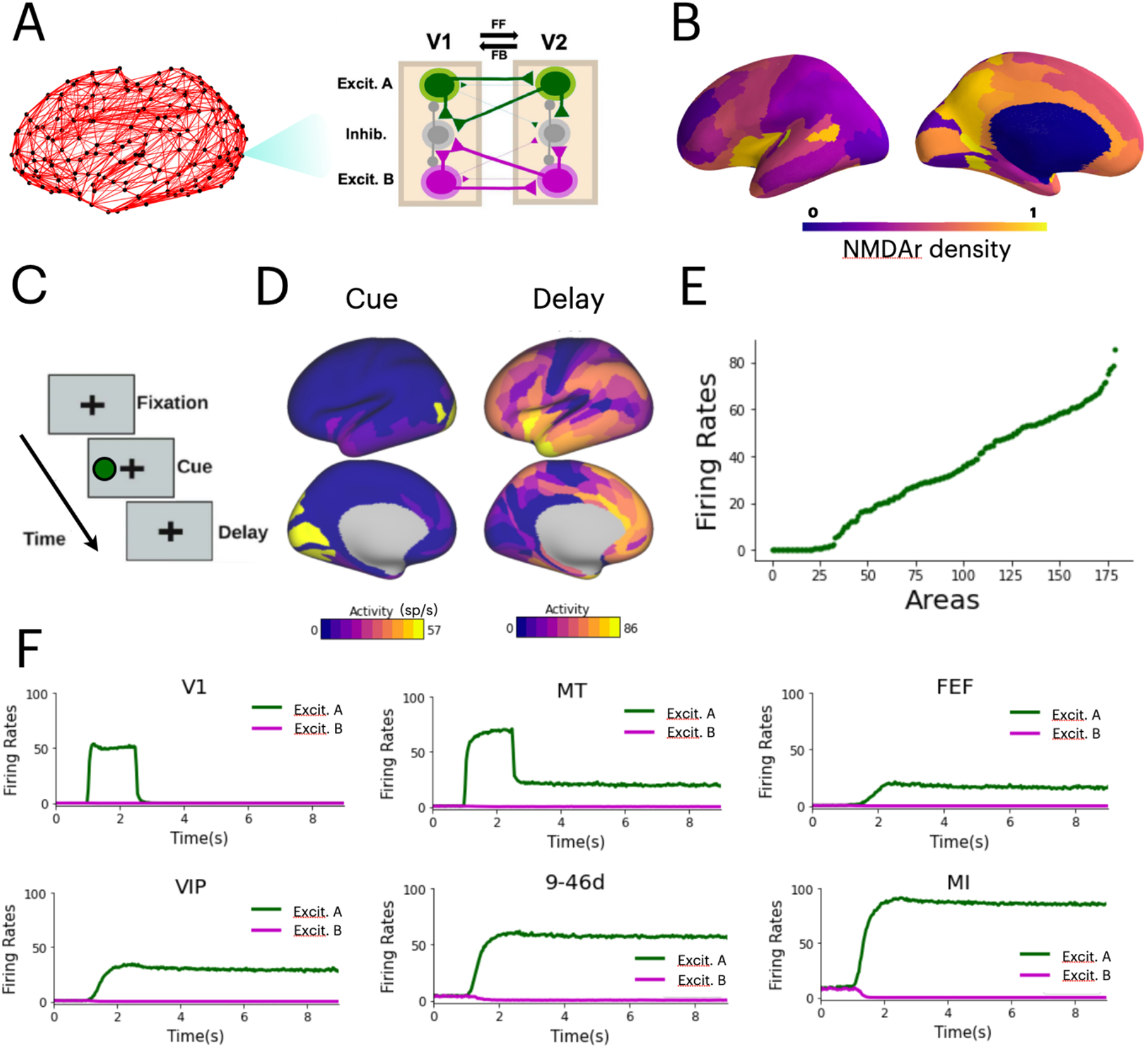
Distributed working memory patterns emerge in a human brain model. (A) Human brain connectome with 180 neocortical areas. Right: each cortical area is modeled as a three population circuit, with two input-selective excitatory populations and one inhibitory population which controls the overall activity of the circuit. Interareal projections to excitatory populations are stronger in the feedforward (e.g. V1 to V2) direction, and those to inhibitory populations are stronger in the feedback (e.g. V2 to V1) direction. (B) Cortical maps (left: lateral view, right: medial view) with the relative density of NMDA receptors in each area. (C) Simplified working memory task simulated here, in which the system has to remember one of two possible visual stimuli (a green circle in this example) after it has been removed from the visual field. (D) Brain activity maps showing the firing rate levels (color code) during the presentation of the stimulus (left) and the delay period (right); the latter displays a broad activation pattern across all lobes. (E) Firing rates for each area in the model during the delay period, ranked as a function of activity. (F) Temporal evolution of selective firing rates (population A in green, B in purple) for several cortical areas. A visual stimulus, for which the excitatory population A is selective, is received for 0.5 seconds.

Most human brain models do not include directional connections in their network, due to the lack of directionality in the MRI-based connectivity used to constrain these models. Recent work^36^ has suggested that the intracortical myelin content of an area (measured by the T1w:T2w map, Fig. S1B) correlates with its position across the cortical hierarchy, thus providing a way to infer directionality from sensory to association areas. Here, we introduced a certain level of directionality in the inter-areal projections of the model by assuming that the hierarchical relationship between two areas modulates the targets in each direction. More precisely, we assume that the feedforward projections, or projections ascending the hierarchy (i.e. from V1 to V2) reach predominantly the excitatory populations in the target area. Likewise, we assumed that projections in the feedback direction (i.e. V2 to V1) were slightly biases towards targeting inhibitory populations (Fig. 1A). This was necessary to ensure that external signals were able to travel from sensory areas up to association areas without significant dissipation, and to avoid overexcitation in the network due to recurrent feedback loops. The bias of the feedforward/feedback projections towards excitation/inhibition can be adjusted with a model parameter, the relative strength of feedback projections targeting inhibitory populations (α). Finally, to introduce a realistic level of heterogeneity in the network, each area has specific parameter values derived from existing data –in particular, the area-specific density of NMDA receptors or NMDAr^37^ (Fig. 1B). The NMDAr map provides information which can’t be inferred from other data sources such as the T1w:T2w map (Fig. S1C,D).

### Distributed working memory emerges in human brain networks

Our first aim was to investigate whether the computational model sketched above was able to replicate the electrophysiological and behavioral features observed in the human brain during a standard working memory task –more precisely, the persistent co-activation of multiple brain areas to selectively retain stimulus-specific information over time after stimulus presentation^21^. We consider a simple working memory task in which the network is presented with a cue, or specific visual stimulus during one second, and after this stimulus is removed, the network has to maintain its identity during a certain delay period (Fig. 1C). The cue is selective to either excitatory population A or B in early visual cortex, which includes area V1 and other secondary visual areas (see Methods for details). During the presentation of the visual cue, the main response of the network is an elevated activity in the visual areas receiving the input (Fig. 1D, left panel). However, during the delay period the activity pattern becomes much broader, with elevated firing being distributed across many cortical areas in different degrees, mostly from frontal and temporal cortices but also from parietal and occipital ones (Fig. 1D, right panel). This emergence of distributed working memory activity is not caused in the model by a local attractor dynamics, as in classical working memory models^39,40^, but rather by long-range interactions between areas which support each other in maintaining the selective activity pattern. Such distributed, self-sustained pattern is in agreement with recent views on distributed working memory patterns in the human brain^21^.

As also suggested by experimental evidence^22^, activity patterns linked to working memory are not fully distributed: some areas are not actively involved in the generation and maintenance of distributed patterns of firing rates. This is reflected in the model by a nonlinear activity profile across all cortical areas (Fig. 1E), in which a good number of areas (~30) maintain a spontaneous level of firing rate (this mostly corresponds to occipital and early sensory areas), and only populations in about 80 areas display persistent firing rates of more than 40 spikes per second. This leads to a very heterogeneous and area-specific dynamics (Fig. 1F): while early sensory areas like V1 react to the visual cue and their activity swiftly decays after stimulus removal, more frontal or prefrontal areas like 9-46d and MI display a persistent pattern of activity. For all areas, the activity is highly selective to the presented stimulus (in this case, stimulus A). We replicated the simulations with a surrogate network in which weights of the inter-areal projections were randomly swapped while maintaining the in- and out-degree connectivity of each area. The emergence of distributed working memory was impaired in this case, demonstrating the importance of the precise large-scale structural brain organization for the existence of the brain-wide activity patterns. (Fig. S2).

### The model explains experimental evidence and predicts forbidden gaps in parameter space

Given that distributed working memory was suggested by our model to be a global phenomenon, we then focused on the two main global model parameters: the overall strength of inter-areal connections or global coupling strength (G), and the relative strength of all inter-areal feedback projections targeting inhibitory populations (α). A first numerical exploration, shown in Fig. 2A, indicates that distributed working memory tends to emerge for large global coupling strengths (indeed no working memory pattern was observed for G<0.6) and moderate levels of feedback inhibition. The maximal value of feedback inhibition permitting the emergence of working memory depends on the global coupling, with higher values of G allowing for stronger inhibitory feedback. Across all the examples depicted in Fig. 2A, the distributed activity patterns remain qualitatively similar to our original case: with the most active areas (in bright colors) located in frontal and temporal areas, and some specific parietal/occipital ones.

**Figure 2:**
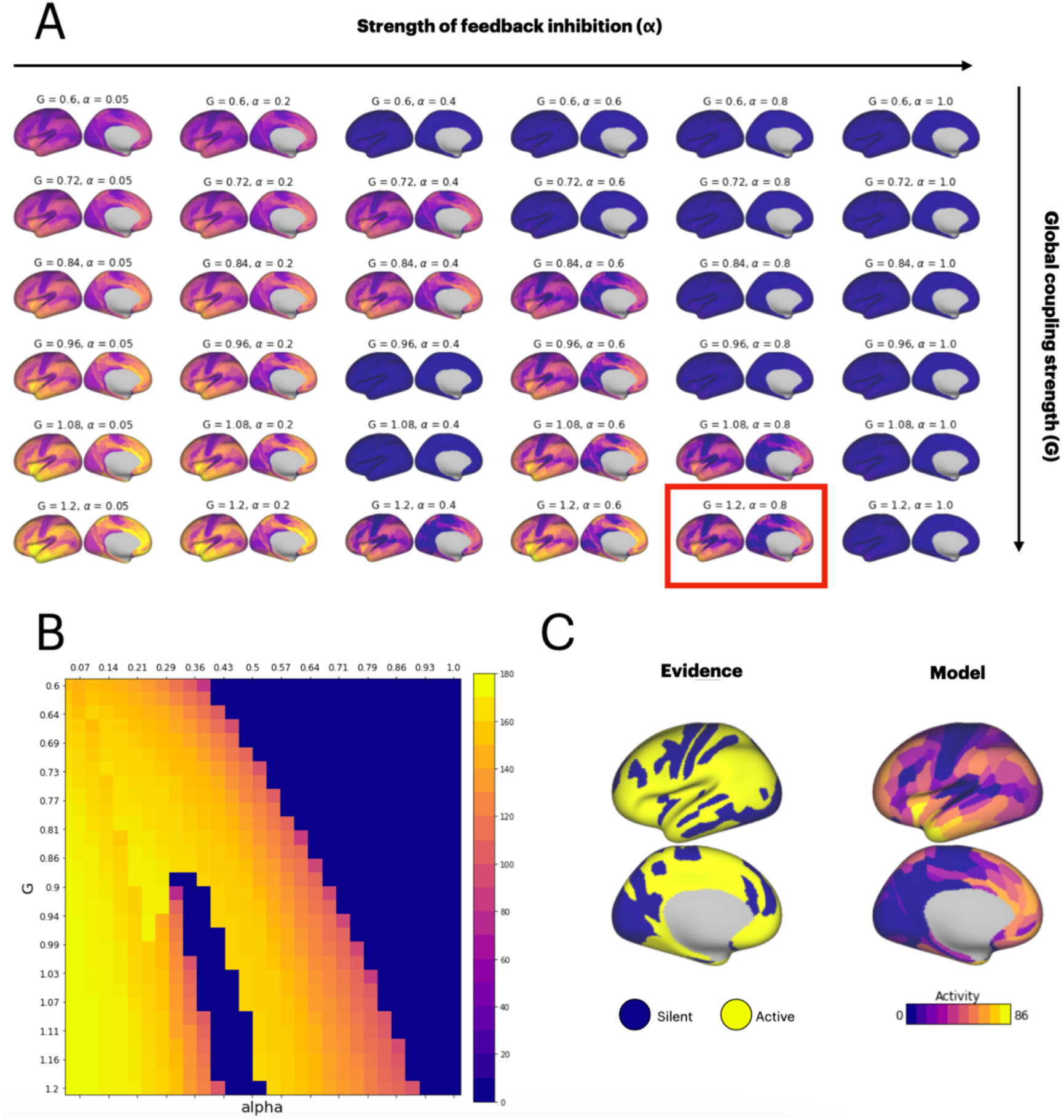
Numerical exploration of the distributed WM states and experimental comparison. (A) Phase space of the model showing the different WM patters obtained as a function of the global couping strenght, G, and the strength of feedback inhibition, alpha. (B) Extensive exploration of the G-alpha space, detailing the existence of a working memory gap for strong global coupling and intermediate strengths of feedback inhibition. (C) Comparison between a cortical map showing areas with sufficient experimental evidence of activity related to working memory (yellow areas on the left map) and the best fit from our model (right), which corresponds to the values of G and alpha framed in panel A.

We also discovered the existence of a forbidden ‘gap’ region in the parameter space, around α=0.4 and G between 0.9 and 1.1, for which no distributed working memory patterns were triggered by the visual input. To ensure that this was not a spurious result of specific simulations, we computed a map of our G-α space with finer detail, showing the overall activity of the network in color code (Fig. 2B). This forbidden gap then was revealed as a localized but extended areas in the center-bottom part of the map. This extended simulation also corroborated the shape of the outer frontier between the working memory phase (bring colors) and silent phase (dark blue, top right side of the map) of our system.

To investigate the factors which originate this forbidden gap, we simulated and alternative version of our model in which the area-specific synaptic strength was given not by the NMDAr density, but by the position of each area in a MRI-estimated cortical hierarchy^36^. Such alternative approach provides a more indirect estimation of local synaptic strength (as it is based on global considerations such as hierarchical position rather than specific local measurements of receptor densities), and the resulting gradient of local synaptic strength becomes more linear and less noisy (Fig. S3). Our simulations revealed (Fig. S3A,B) that this alternative version of the model did not have a forbidden gap area, and that the origin of such gap in our original model is the large level of heterogeneity introduced by NMDAr density data. Indeed, when the heterogeneity of NMDAr is reduced to a linearized gradient as in the MRI-estimated hierarchy (Fig. S3C,D), the forbidden gap progressively closes (Fig. S3E,F). Without the accurate, area-specific data on NMDAr densities, our human brain model would have been unable to predict the existence of the forbidden gap.

To quantitatively compare our model predictions with experimental evidence, we performed a preliminary literature search (Table S2) to identify the cortical areas most likely related to working memory maintenance, in a similar way to recent revisions for macaque^22^. We then compared the resulting map of experimental evidence to all model results (with thresholded activity) in our G-α space, and identified the parameter values which provided the best fit. The comparison between the evidence map and the best model fit (Fig. 2C) yielded a degree of overlap of about 78% –in other words, the large majority of brain areas participating in distributed working memory patterns can be explained with our model by simply fitting two parameters (G and α). The parameter values providing the best fit (highlighted in Fig. 2A with a red frame) correspond to a strongly connected network with moderately strong feedback inhibition. Other metrics provide also high levels of overlap with the data (Fig. S4).

### Robustness to interferences from other sensory modalities and distractors

So far, we have only considered a simple task containing the minimal ingredients needed to explore working memory: a brief cue presentation and a delay period. Tasks used in experimental conditions are far more complicated and contain more complex elements. To test whether our distributed working memory is resilient to other task elements, we introduced a GO signal at the end of our delay period, similarly as how experimentalists indicate macaques in real working memory experiments that the delay period is over and an action has to be taken. To also test the impact of introducing other sensory modalities in the process, we considered that this GO signal was a non-selective auditory cue (i.e. the auditory signal activated both excitatory populations of the primary auditory areas, Fig. 3A).

**Figure 3:**
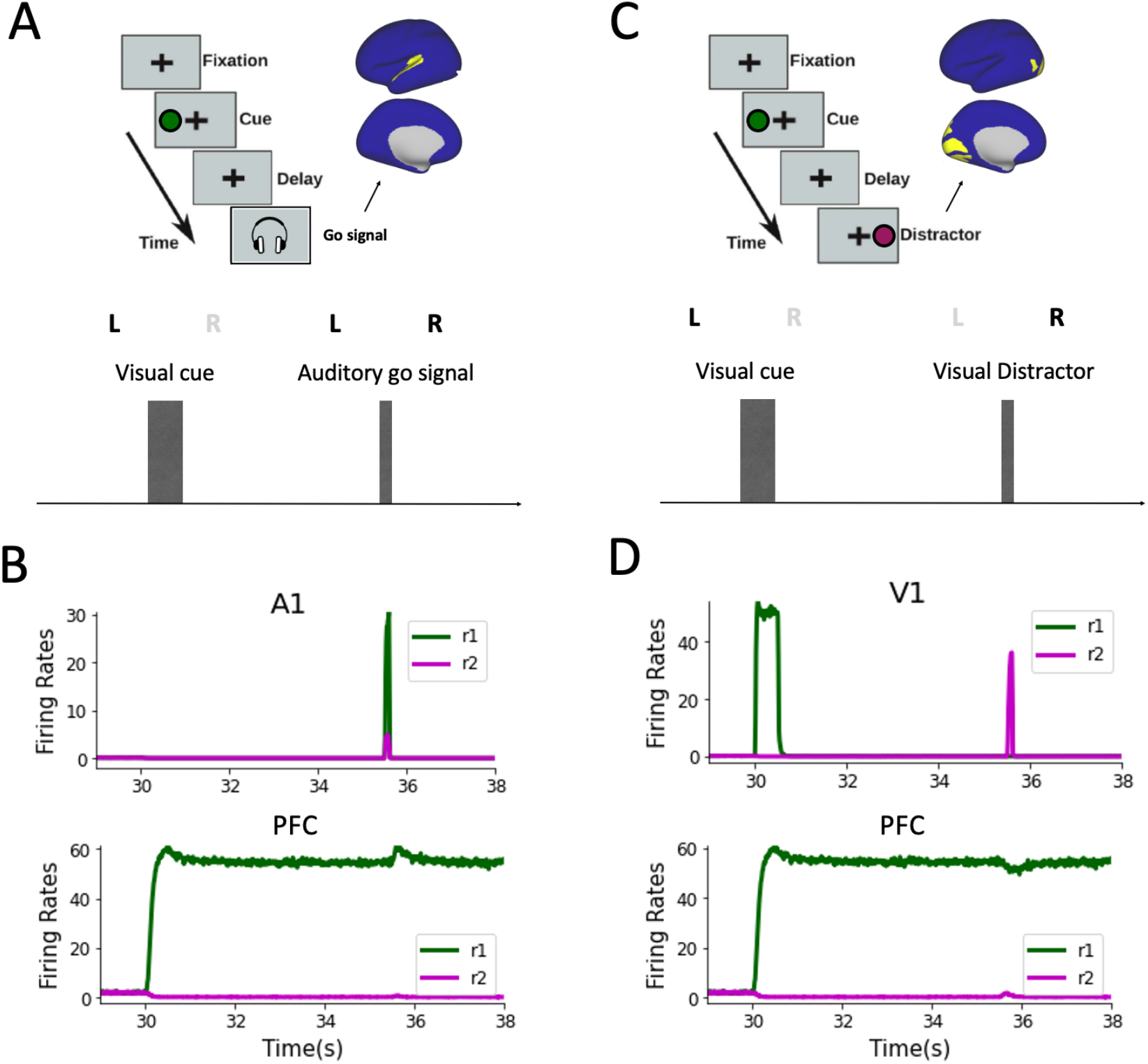
More complex working memory tasks: auditory GO signals and visual distractors. (A) Delayed task with an auditory, unspecific GO signal. Brain map show areas affected by the auditory GO signal in yellow. (B) Response of primary auditory area A1 and prefrontal cortex (area PGS), showing the robustness of the working memory state to unspecific signals from other modalities. (C) Delayed task with an opposing visual distractor (purple circle). Brain maps show areas affected by the visual distractor in yellow. (D) Response of primary visual area V1 and prefrontal cortex (PGS), displaying robustness of the memory to selective visual distractors. Both the auditory GO signal and the visual distractor have the same strength as the visual cue, although a shorter time duration.

While the GO signal triggered a strong response in both excitatory populations of primary auditory area A1, the signal did not disrupt the encoding of pattern A previously specified by the visual cue (Fig. 3B), leading only to a small bump in the activity of prefrontal areas like PGS. This reflects the robustness of the distributed encoding to other potentially disrupting sensory modalities. Similarly, we investigated the robustness of visually-cued distributed representations after an opposite visual signal, or distractor, was presented during the delay period (Fig. 3C). The distractor was of the same intensity as the original visual cue, but of shorter duration. We found that the distractor was not able to disrupt the selective working memory encoded in higher areas, including prefrontal cortical areas like PGS, where only a small dip in firing rate was observed (Fig. 3D). As in the case of computational models of NHPs^32^, the robustness of the memory pattern was due to a combination of the distributed nature of the attractor and the inhibitory feedback provided by active frontal/prefrontal areas to sensory areas, which notably decreased the impact of subsequent stimuli when an opposing item is already stored in memory.

### Comparison between human and NHP models

Driven by the partial agreement between our human brain model and previous models of distributed working memory in NHPs, as for instance in terms of resistance to distractors (Fig. 3D), we then replicated a large-scale macaque model of distributed working memory^32^ and compared its results with our human brain model, to perform an in-silico comparison of distributed working memory across species. Despite both models presenting severe structural differences (for example, in the number of cortical areas, connectivity patterns, projection directionality, and other aspects), it is possible to make an approximate comparison between visually evoked distributed memory patterns in both models. In both human and NHP models the set of cortical areas active during working memory is fairly distributed across most of the cortical surface (Fig. 4A), with the highest activity located in temporal and frontal lobes in both cases. When normalized with respect to the highest activity on each lobe, however, we observe significant differences between human and NHP activity patterns. For example, normalized activity levels in frontal and parietal areas tends to be significantly lower in the human brain model than in the NHP brain model (Fig. 4B). No significant differences were found however between prefrontal, temporal or occipital lobes of both models.

**Figure 4:**
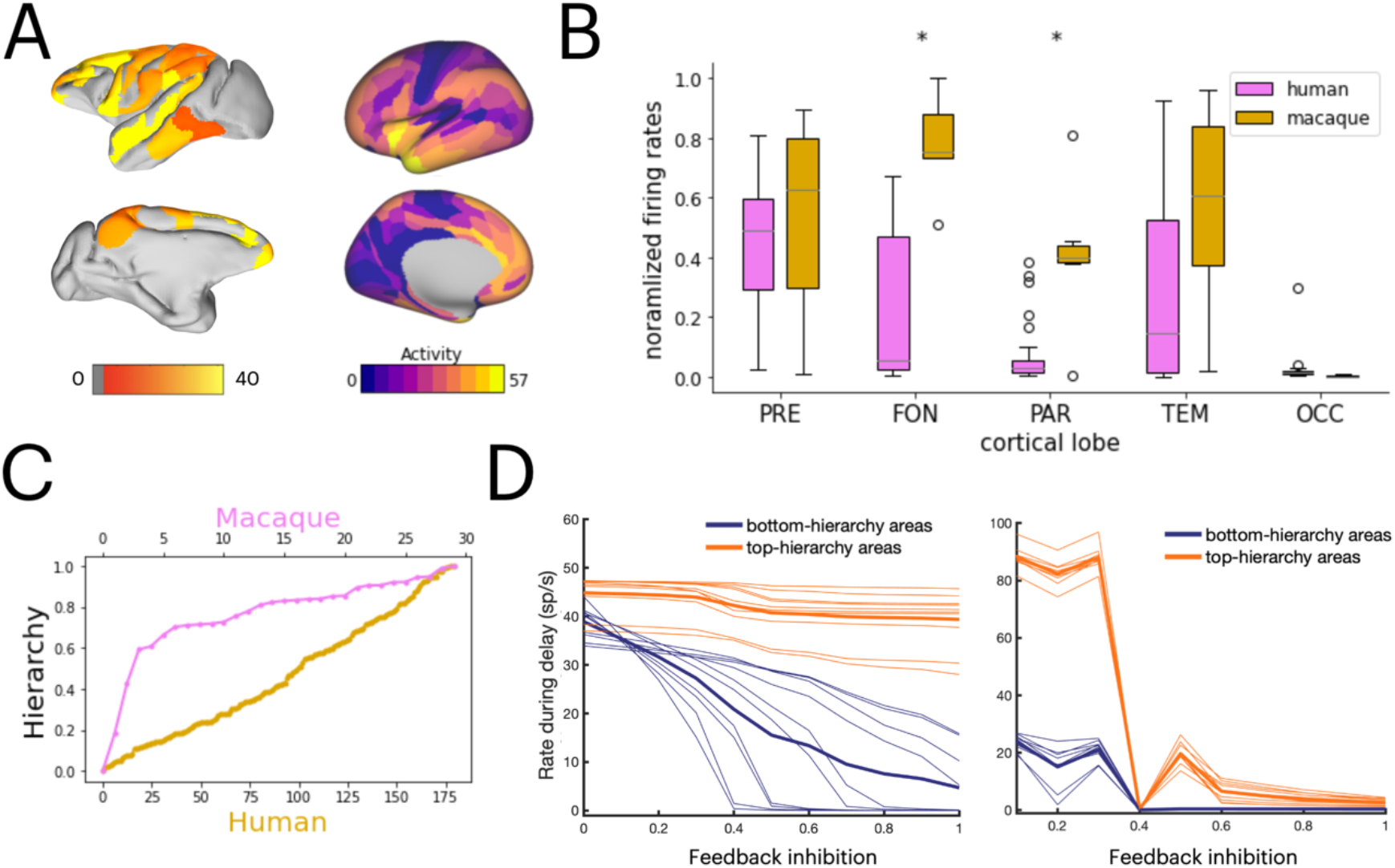
Comparison between distributed working memory models for human and NHP. (A) Brain maps showing distributed activity during the delay period for a macaque brain model (left) and a human brain model (right). (B) Normalized averaged firing rates for prefrontal, frontal, parietal, temporal and occipital areas, for both models. Differences in frontal and parietal lobes are significant for P-FDR<0.05. (C) Normalized local coupling strength across the hierarchy for both models, displaying the differences in shape and slope. (D) Effects of varying the strength of feedback inhibition on the average activity of top-hierarchy (orange) and bottom-hierarchy (blue) areas for the NHP (left) and human model (right).

One of the key structural differences between both models was how areas are distributed across the cortical hierarchy. In the NHP model, hierarchical distances between areas followed estimations based on tract-tracing data^41,42^, which yield steep hierarchical relationships between early visual areas and a reduced slope for associative areas (Fig. 4C). A consequence of this is that a strong feedback inhibition from associative areas to sensory areas (equivalent to our parameter alpha in the human model) was needed to prevent a full-brain activation. Indeed, we observed a clear distinction between delay firing rates between top- and bottom-hierarchy areas only when the feedback inhibition is sufficiently strong (Fig. 4D, left panel). In the human model, however, the MRI-based cortical hierarchy^36^ leads to a much more linear hierarchy (Fig. 4C). In this case, feedback inhibition was not drastically needed to obtain the partially distributed patterns suggested by experimental evidence. A clear distinction between in top- and bottom-hierarchy firing rates was observed for any values of feedback inhibition –except for α~0.4, which corresponds to the forbidden gap (Fig. 4D, right panel).

### Distributed activity patterns explain the decoding of memory contents from V1 activity

While substantial experimental evidence exists for activity-related working memory in prefrontal and, to a lesser extent, temporal and parietal cortex, the involvement of occipital cortex is still under debate^21,22,43–46^. Since our data-constrained model allows to explore the role of any cortical area in working memory, it is uniquely well positioned to study how working memory contents may be decoded from activity in early sensory areas such as V1^43,44,46^. We built a linear classifier to decode which memory item (pattern A or B) is being held in a distributed working memory pattern, and applied it to the firing rates of primary visual area V1. For this analysis, we sampled a restricted range of values in the G-α space (Fig. 5A, top panel), and stimulated the system with a brief, selective visual cue (Fig. 5A, bottom panel).

**Figure 5:**
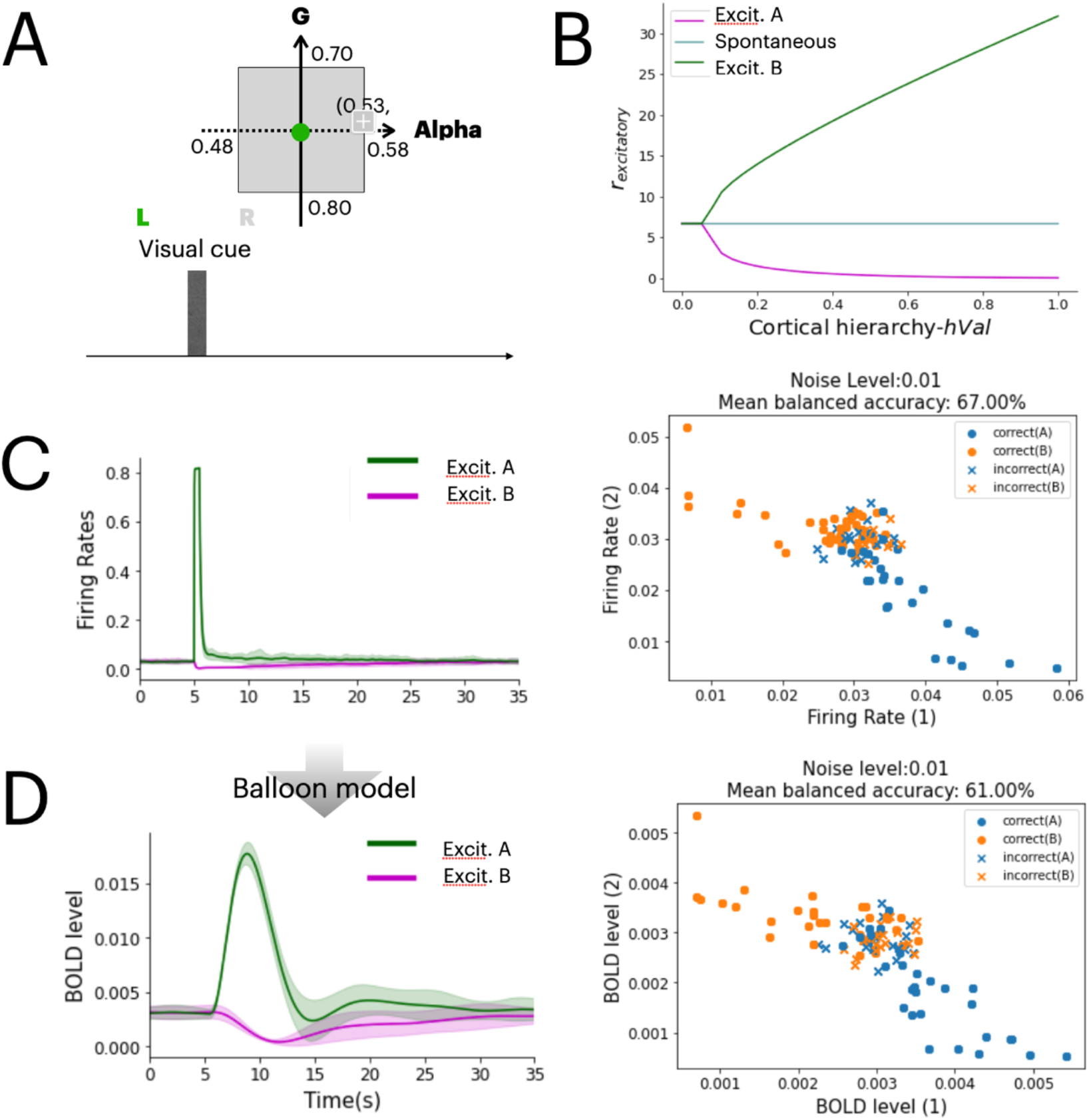
Decoding the contents of working memory from V1 activity. (A) Top panel: range of G and alpha considered for the decoding of V1 activity. Bottom panel: experimental scheme where a visual stimulus on the left is briefly shown. (B) Firing rates for all areas during the delay period for the selected (green) and non-selected (purple) populations, and the spontaneous state (gray). Decoding between stimuli is hardest for early sensory areas (left side). (C) Firing rates of V1 excitatory populations during sensory presentation (shading denotes standard deviation over 50 trials), and decoding of the visual stimuli from firing rates values, for G = 0.75, alpha = 0.53, and a noise level of σ_A,B_= 0.01. (D) Same as panel C, but with BOLD signals obtained from the Balloon-Windkessel model instead of firing rates.

The cue triggered a transient response in V1 and a distributed working memory pattern with selective persistent activity across other areas during the delay (Fig. 5B). While discerning between the firing rates of excitatory populations A and B is simple for prefrontal and other high areas, the task is harder for early sensory areas such as V1, where shortly after stimulus inactivation both excitatory populations have similar activity (Fig. 5C, left panel). Despite this, the linear classifier was still able to decode which pattern was being held in memory from V1 firing rates, with an accuracy of 67% (Fig. 5C, right panel). The decoding accuracy was defined as the number of trials with a successful decoding divided by the total number of trials. The model indicated that this successful decoding was possible due to the long-range interactions between higher association areas, in which memory-selective activity was more marked, and V1, which was still able to reflect subtle differences despite receiving strong feedback inhibition. To better replicate the decoding done in experimental conditions, we used the Ballon-Windkessel model^47^ to transform the conductance variables of the model into BOLD signals (Fig. 5D, left panel) and repeated the process. The decoding from BOLD signals in V1 was able to predict the stored information with an accuracy of 61% (Fig. 5D, right panel). Other parameter values of G and α also lead to high classification accuracy (Fig. S5A,B). The decoding accuracy was also reduced by the level of noise and the strength of feedback inhibition, but increased with the global coupling strength (Fig. S5C).

## Discussion

Computational models of the human brain are a versatile and valuable tool to understand neural brain dynamics, although the connection between such brain dynamics and mechanistic implementations of cognitive functions has been elusive. Aided by recent experimental and computational findings, we presented in this work a first example of a computational model which introduces a cognitive function such as working memory in a full-brain dynamics framework. Our achievement was possible due to the combination of three specific datasets: a human connectome^34,35^, a cortical hierarchy estimate^35,36^ and a NMDAr density map^37^. Distributed working memory patterns were able to emerge in such a simplified network model, with the stimulus-selective elevated activity being sustained by long-range interactions between brain areas rather than by local recurrent dynamics as assumed by classical attractor models^39,40^. The model has also able to explain a large fraction of the experimental variability for working memory-related activity in the brain, and provided predictions regarding forbidden gaps in the parameter space, inter-species comparisons and content-decoding in sensory areas.

The distributed nature of working memory is an aspect which has caught much attention in recent years. Detailed review for human and NHPs^21,22^ provided an overview of the many areas potentially involved in working memory, but didn’t deliver a mechanistic insight as of how such distributed activity patterns would emerge and what were the neurophysiological implications. Recent experimental efforts have given more conclusive evidence of the distributed working memory patterns for the case of mice^25^, triggering further interest on whether working memory would display similar traits in the case of human and other primates. On the other hand, computational modeling work has explored the idea of distributed working memory in macaques across various levels of detail along recent years^23,48–50^, with the first complete computational study of distributed working memory in a data-constrained macaque network recently proposing the core mechanistic principles of the phenomenon^32^. While similar efforts have been carried out for the case of mice^33^, the case of distributed working memory in humans had not been tackled until now.

One of the main ingredients of the model is the consideration of area-specific heterogeneity. This has been achieved by constraining the synaptic efficacies via PET-derived, area-specific density of NMDA receptors^37^. PET-derived receptor densities provide a convenient proxy to estimate the synaptic strengths in a particular brain area, and the data used here has also been successfully validated by comparing it with more precise autoradiography data of NMDAr density^51,52^. To consider proper directionality in the connectivity –a common limitation of human brain computational models since neuroimaging-derived structural connectomes have non-directional projections –we introduced in our model an MRI-derived estimation of the hierarchical ordering between all 180 areas^36^. While this hierarchy is not as anatomically precise as the ones obtained for NHPs via tract-tracing techniques^41,42,53^, it provides a first estimation of directional interactions which is strongly correlated with the ground-truth cortical hierarchies in NHPs^36^. This hierarchical architecture offers a structural ground for the temporal functional hierarchies observed in human neuroimaging studies^54–59^, similarly to the case of temporal spike cross-correlations in macaques^32,60,61^.

In a first attempt to compare with experimental evidence of distributed working memory in the human brain, we have gathered information about those areas most likely involved in working memory in humans, and collected an ‘evidence list’ (see Table S2) similar to the ones proposed for macaque^22^. One should not expect of course to find all these areas simultaneously co-activated in any given working memory task, since this evidence map was obtained by aggregating results from studies using many different tasks. The comparison with our model results is however appropriate, since the model doesn’t include any specific details to simulate advanced or specific working memory tasks –beyond the elemental examples explores in Fig. 1 and 3 –and generated therefore a ‘maximal’ distributed attractor state which we compared to the map of evidence aggregated across tasks, with a degree of overlap of about 80% between model and data. To successfully reproduce the particular distributed pattern of, for example, a numerosity working memory task, the sensitivity or specificity of neural activity of each cortical area for the task at hand should be implemented in the model, something which has been previously tested for macaque models^32^ and could make use of any normalization of gating mechanism at the neuron or circuit level^49,62–66^.

While several of the conclusions from our model are shared with the ones obtained for the case of macaques and mice (such as the key role played by long-range projections in the emergence of distributed working memory patterns), the model indicates multiple key differences between the way that distributed working memory emerges in human and other animals. For example, the assumption of having a strong bias of feedback projections towards targeting inhibitory neurons is necessary in the macaque model to achieve realistic low-activity levels in sensory areas^32^ (see also Fig 4D) and to replicate multiple experimental evidence across scales^67^. However, those low-activity levels can be obtained in the human model with less restrictive conditions (although strong feedback inhibition provides the best fit with the data). This suggests that the linearity of hierarchical interactions between human brain areas eliminates those restrictions for the human brain, providing additional flexibility in the way top-down projections might modulate perception in the case of humans. Differences with the case of mice are even more salient: while current efforts identify gradients of inhibitory neuron density as a fundamental factor for the distributed patterns^33^, our model suggests that variations in NMDAr densities are enough to explain the evidence in humans. More detailed models will allow for a deep computational study of the differences in working memory mechanisms across these and other species.

The present model allows, for the first time, to integrate realistic full-brain dynamics with mechanistic functionalities linked to a cognitive function. This opens the door to address many important questions requiring both a functioning architecture for cognitive functions and a realistic neural dynamics across whole-brain networks. For example, our model allowed us to explore the activity profiles of early visual areas during working memory tasks in a way that other human brain models have been unable to do. The model revealed that the influence of long-range feedback projections from prefrontal cortex to early visual areas was enough to allow the successful decoding of working memory contents from BOLD signals recorded in V1, as observed experimentally. The fact that this feedback influence was mostly inhibitory did not prevent a successful decoding, however the maintenance of the selective information occurs primarily within a distributed networks of frontal and prefrontal areas. While V1 is vital for propagating the visual signal early in the trial, suppressing its activity does not disrupt the information stored in working memory in a significant manner, supporting the theory that V1 may reflect working memory contents but it is not vital for its maintenance^43–46^.

In the current model, we considered that working memory was maintained via selective persistent activity underlying attractor dynamics, in line with traditional approaches^39,40^. However our main principle for distributed working memory is compatible with other mechanisms, such as time-varying activity mechanisms^16–18,20,68^ or silent working memory^14,15,18,69^, as computationally tested for macaques^32,50^. The role of thalamic nuclei in working memory maintenance^23^ aligns well with our core proposal, and the addition of thalamic subnetworks to our model would arguably lead to a more robust working memory mechanisms, also in line with previous modeling efforts of cortico-pulvinar systems^49^.

The present study provides three well defined predictions suitable to experimental testing. First and foremost, our model predicts that activity-related working memory relies not only on local recurrent interactions, but also on long-range communication between distant brain areas. Therefore, disrupting or modulating the activity of areas beyond prefrontal cortex (for example, using transcranial magnetic stimulation) should impact working memory performance. Second, we predict the existence of a ‘forbidden gap’, or a certain set of values of the feedback inhibition strength (assuming strong enough G) which makes impossible to sustain working memory activity. Pharmacological approaches able to diminish or partially block receptor activity in middle and early sensory areas could be able to detect such gaps. Third, we predict that functional connectivity might reflect a brain-wide distribution of interactions during the delay period of a working memory task, in contrast to the resting state case in which structural and functional connectivity will be more similar to each other. Fourth, the model indicates that decoding working memory contents from V1 is possible, but inactivation of V1 should not destabilize working memory maintenance.

In conclusion, we present the first data-constrained human brain model able to mechanistically simulate a cognitive function, and we show its level of agreement with available evidence. We foresee that this new type of models, which combine realistic large-scale brain dynamics with cognitive capabilities, will play a fundamental role in our understanding of distributed cognitive process in humans, and uncover potential origins of their malfunction when coupled with tools of computational psychiatry.

## Acknowledgments

We thank Nicola Palomero-Gallagher for insight in the PET-derived data and Jesús Cortés for his valuable feedback on an early version of the manuscript.

## Funding

This work was supported by the Human Brain Project Flagship project 945539 (JFM) and a UvA/ABC Project Grant (JFM).

## Author contributions

JFM conceived the study; MF, AB and JFM designed the computational experiments, performed the research, analyzed the results, and wrote the manuscript.

## Competing interests

Authors declare not competing interests.

## Data and materials availability

All information needed to reproduce the results of this manuscript are in the main text and Materials and Methods section, and the code used will be made available upon publication of this work.

## Methods

### Computational model: local circuit

We start by describing the dynamics of a typical local cortical microcircuit, representing a brain region, via the Wong-Wang model^38^. The version that we use here simulates the temporal evolution of the firing rate of two interconnected, input-selective excitatory populations, as well as the evolution of the firing rate of an inhibitory population also interconnected with both excitatory populations (see Figure 1A). The dynamics is given by the following equations:

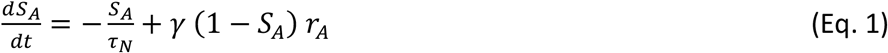

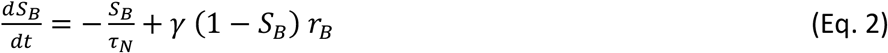

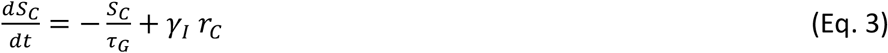

The variables *S_A_*, *S_B_* and *S_C_* represent the NMDA conductances of both selective excitatory populations and the GABA conductance of the inhibitory population respectively. The time constants are *τ_N_* = 60 ms*,τ_G_* = 5 ms*,γ* = 1.282 and *γ_I_* = 2. The variables *r_A_, r_B_* and *r_C_* are the mean firing rates of the corresponding excitatory and inhibitory populations, which evolve in time as follows:

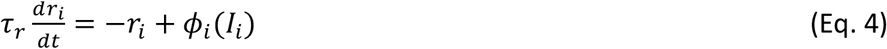

where the time constant for the firing rates is τ_r_=2 ms, and the so-called transfer functions for the excitatory and inhibitory populations are given by:

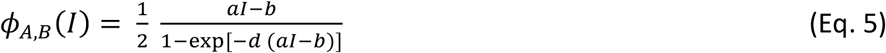

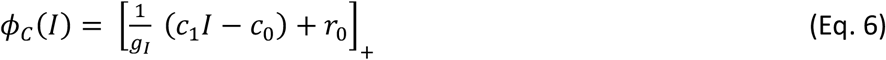

The values for the parameters are *a*=135 Hz/nA, *b*=54 Hz and *d*=0.308 s for the excitatory populations, and g_I_=4, c_1_=615 Hz/nA, c_0_=177 Hz and r_0_=5.5 Hz for the inhibitory one. The notation [*x*]_+_ denotes rectification.

The input current I_i_ for each population ‘i’ is given by

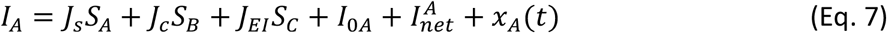

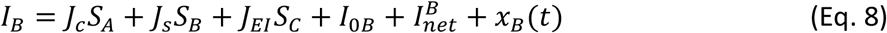

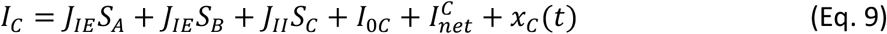

In these equations, J_s_, J_c_ are the self- and cross-coupling between excitatory populations. Likewise, J_EI_, J_IE_ and J_II_ are, respectively, the coupling from the inhibitory populations to any of the excitatory ones, the coupling from any of the excitatory populations to the inhibitory one, and the self-coupling strength of the inhibitory population. The parameters I_0i_ with i=A, B, C are background inputs to each population. Their values are J_s_=0.3213 nA, J_c_=0.0107 nA, J_IE_=0.15 nA, J_EI_=−0.31 nA, J_II_=−0.12 nA, I_0A_=I_0B_=0.3294 nA and I_0C_=0.26 nA.

Some of the parameters above, in particular J_s_ and J_IE_, will be modified in the section below in an area-specific way to introduce a gradient of properties across the cortical hierarchy. Sensory stimulation can also be introduced here as extra pulse currents of strength I_pulse_ =0.3 and duration T_pulse_ =0.5 sec (unless specified otherwise). Finally, the term I^i^_net_ denotes the long-range input coming from other areas in the network (which we assume zero for now), and the term x_i_(t) with i=A, B, C is an Ornstein-Uhlenbeck noise process, given by

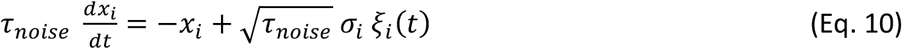

ξ_i_(t) is a Gaussian white noise, the time constant is τ_noise_=2 ms and the noise strength is σ_A,B_=0.005 nA for excitatory populations and σ_C_=0 for the inhibitory one.

### Computational model: gradient of cortical properties

Local cortical circuits may display important differences across cortical areas, and such differences might significantly impact the contribution of each area to functions like working memory. Here, we introduce area-level heterogeneity by adjusting the synaptic weights of each area, following recent positron emission tomography data which estimates the density of NMDA receptors (or NMDAr) per area in the human brain^37^ (see Fig. 1C). For that, we mapped their 34-area parcellation of the NMDA receptor density profile to the 180 area parcellation for the simulation. The NMDAr density serves in the model as a proxy for the number of synaptic terminals in each area, and therefore also for the overall strength of macroscopic synaptic interactions.

In particular, we introduce the cortical heterogeneity in the model by assuming that the synaptic strength *J_s_* has area-specific values which are proportional to the NMDAr density of each area d^i^_N_, as follows:

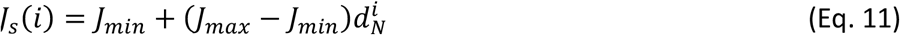

Here, we set *J_min_*=0.205 nA and J_max_=0.42 nA, unless specified otherwise. For reference, it is important to note that an isolated local circuit like the one specified here won’t be able to display bistability unless J_max_ is set above 0.4655 (at which point a bifurcation from monostable from multistable dynamics occurs^32^). Therefore, at the present values of J_min_ and J_max_, all persistent activity observed in the system will be a consequence of global interactions between cortical areas, instead of the result of local bistability as in classical attractor models of working memory.

To maintain the excitatory input at the spontaneous level similar for all areas (and therefore have similar *spontaneous* firing rates across all areas and within physiologically realistic values), an increase of J_s_ for a given area needs to be compensated by a local increase in inhibition. To ensure this, we follow the same strategy as in a previous study^32^ and assume a linear relationship between the synaptic strength from excitatory populations to the inhibitory population (*J_IE_*) and the recurrent excitatory synaptic strength (J_*S*_), given by

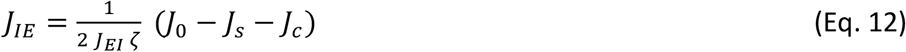

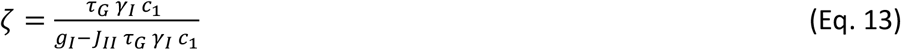

As an alternative take on the issue of area-level heterogeneity, specifically explored in Supplementary Fig. 3-1, we also assume a variation of our model in which area-specific synaptic strengths are given by the position of each area in the so-called cortical hierarchy, instead of by each area’s NMDAr density. Previous work in macaques has demonstrated that the position of an area within the anatomical hierarchy^41,42,53^ positively correlates with the number of dendritic spines per pyramidal neuron^32,61^, which serves as a proxy for synaptic density. While a traditional anatomical cortical hierarchy as in Felleman and Van Essen^53^ is not available for humans, recent studies have suggested that MRI-derived T1-to T2-weighted mapping (T1w:T2w) may serve as a suitable alternative for humans^36^. For this alternative version of our human brain model, explored in Fig. S3, the self-coupling synaptic strength is given by

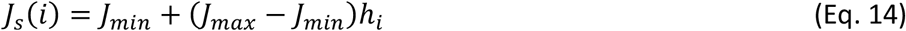

Here, h_i_ is the position of area ‘i’ in the T1w:T2w ranking given in previous work^36^.

### Computational model: interareal projections

For interareal connections, excitatory populations (A or B) in a target area receive feedforward projections from the correspondent excitatory population (A or B) for each source area projecting to it. Feedforward projections have a relative strength F^xy^, with ‘y’ being the source area and ‘x’ being the target area. Likewise, the inhibitory population (C) receives feedback projections (of relative strength 1-F^xy^) from both excitatory populations of such source areas.

The total sum of the interareal current received by a population A/B/C in area ‘x’ is given by

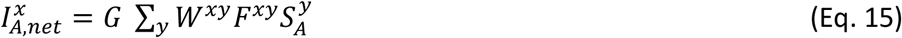

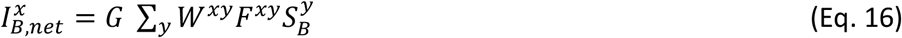

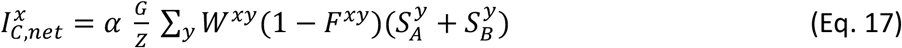

The first factors are the global coupling strength G (=0.48 unless specified otherwise), the relative strength of (feedback) inhibitory projections α, and the excitatory-inhibitory balancing factor Z, which guarantees an approximate cancelation of excitatory and inhibitory effects at spontaneous activity levels, as in previous studies^32^. It is given by

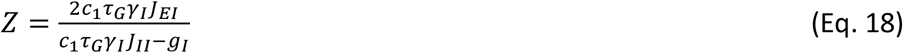

The second relevant factor in Eqs. 15-17 include the connectivity strength from source area ‘y’ to target area ‘x’, or *W^xy^*, which can be obtained from the original connectivity weights from diffusion imaging (D^xy^) by range compression with parameters k_1_=1.2 and k_2_=0.3:

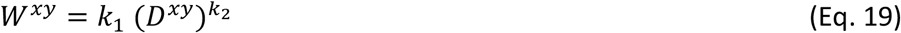

Finally, feedforward (F^xy^) and feedback (1-F^xy^) factors were derived from the T1w:T2w ratio provided by the MRI-estimated hierarchy^36^. Specifically, we assume that (i) the F^xy^ strength from hierarchically horizontal connections (i.e. when x=y) is 0.5, and (ii) as the hierarchical distance between target and source areas increases, the F^xy^ strength increases (decreases) if the source area is lower (higher) in the hierarchy than the target area. The relationship between F^xy^ and the data-derived hierarchical distance is piecewise: when the source area moves up in the hierarchy from 0.5, F^xy^ decreases (increases) with hierarchical distance, leading to increasing areas with F^xy^ values of zero (one). The relationship is opposite when the source area moves down in hierarchy from 0.5. Intuitively, this means that when an area moves up in the hierarchy, it sends increasingly less feedforward inputs to target areas.

### Simulations and analysis

#### Sensory input

We model the external visual input as targeting a small set of early visual areas: V1, VM1, V3A, VMV3, V4, MT and V6A. Such choice is motivated by the fact that thalamic projections target visual areas beyond the primary visual cortex V1. The visual input is selectively given to one excitatory population (A or B) in those areas, with an intensity of 0.3 nA for 0.5 second. For the auditory GO-signal, we give inputs (c = 0.3 nA) to both excitatory populations of seven auditory areas: A1, RI, TA2, PBelt, MBelt, LBelt and A4.

#### Working memory tasks

A simple visual delayed response task is used to test the model performance, where a brief (0.5 second) excitatory input is given to a selective neural population (A or B) of the visual areas specified above (Fig. 1C). Estimations of the firing rate levels for each area are performed by averaging the activity of the last two seconds at the end of the delay period. To further evaluate the robustness of the model, we also simulated two more complicated working memory tasks (Fig. 3). For both tasks, the system needs to remember the position (left or right) of a triangle shown on a screen. After a fixation period, the stimulus is received by early visual areas for a brief time (0.5 second) before being removed. After a delay period of five seconds, we introduced (i) a brief (0.1 seconds) binaural auditory stimulus (auditory GO-signal), for the first task, or (ii) a visual distractor of opposite selectivity than the original visual cue (0.1 seconds, c=0.3 nA).

#### Hemodynamic transformation and functional connectivity

To consider in the model the hemodynamic transformation from our conductance variables to a blood-oxygen-level dependent (BOLD) signal, we use the Balloon-Windkessel model^47^, a mechanistic model for the hemodynamic response. Following previous approaches^36^, the equations describing this transformation are:

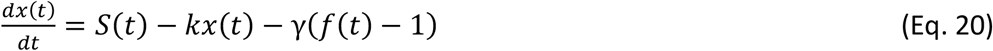

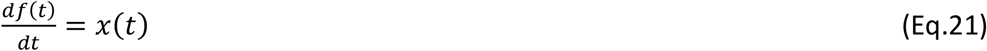

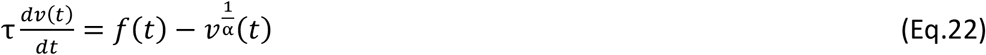

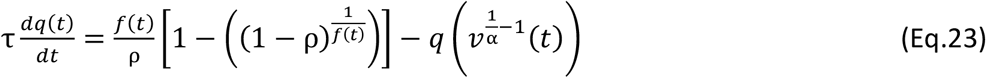

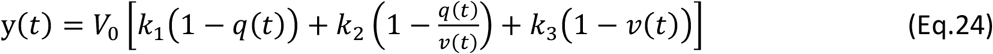

Here, S is an excitatory synaptic gating variable (corresponding to S_A_ or S_B_ in our local model above), x is the corresponding vasodilatatory signal, f is the blood inflow for that population, v is the corresponding blood volume, and q is the deoxyhemoglobin content. The parameter ρ = 0.34 denotes the resting oxygen extraction fraction, τ = 0.98 s is the hemodynamic transit time, and κ = 0.65 Hz is the rate of signal decay. Finally, k_1_= 3.72, k_2_= 0.53, and k_3_=0.53 are dimensionless magnetic field strength-dependent parameter values, γ = 0.41 Hz is the rate of flow-dependent elimination, α = 0.32 is the Grubb’s exponent, and V_0_ = 0.02 is the resting blood volume fraction.

#### Macaque model

The macaque/NHP model used in the comparison with the human working memory model is taken from a previous work^32^. Briefly, it consists of a large-scale network of 30 areas from data-constrained macaque neocortical connectome, with directionality inferred from the position of each area in the cortical hierarchy and local synaptic strength estimated from local spine density data. The local circuit model is identical to the one used here, and inputs go to V1 with same length and strength as in human model.

#### V1 decoding

To test whether visual stimuli can be decoded from neural activity during delay period in V1, we repeated the simple visual delayed response task for 50 times for each stimuli (left or right) with G = 0.75, alpha = 0.53 and noise level = 0.01. We transformed the model outputs to BOLD signals and we then measured the mean BOLD level for both neural population during delay period for each trial. The resulting data is used to train linear classifiers via logistic regression to decode the stimuli. The classifiers were cross-validated with the data and the resulting decoding accuracies (defined as the number of trials with a successful decoding divided by the total number of trials) were recorded. We then varied the values of several parameters to see their effect on decoding accuracy: noise levels ranging from 0 to 0.02, alpha values from 0.48 to 0.58 and G values from 0.7 to 0.8.

---

**Figure S1:**
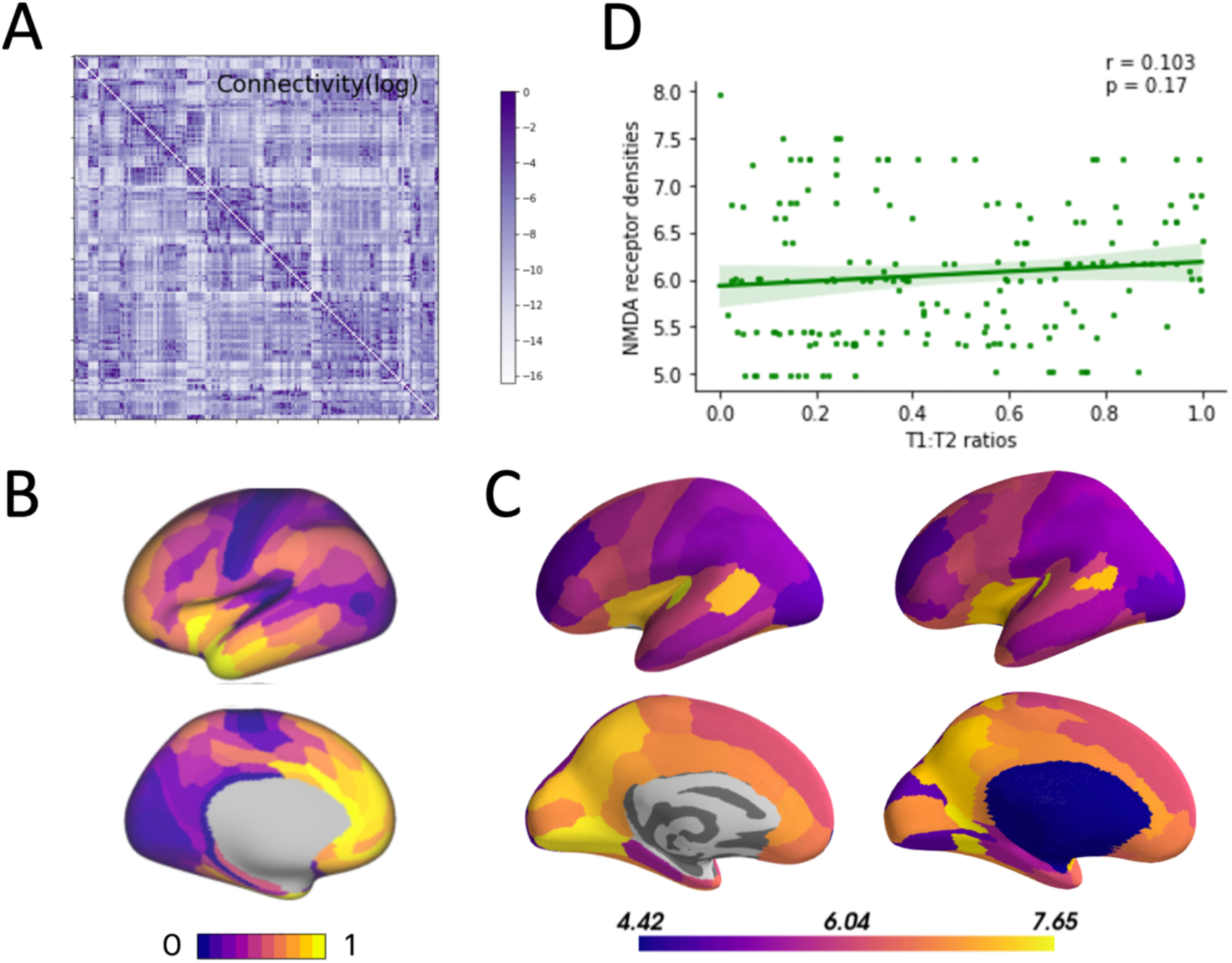
Anatomical data for human model. (A) Human connectivity matrix (180 ROIs, left hemisphere) from MRI data from the Human Connectome Project (see main text for details). (B) MRI-based T1w:T2w map. (C) NMDA receptor density map. Left panel shows the map for parcellation in the original study, and right panel shows the map we generated for the parcellation we are using in the model. (D) Correlation between T1w:T2w values and NMDA receptor density.

**Figure S2:**
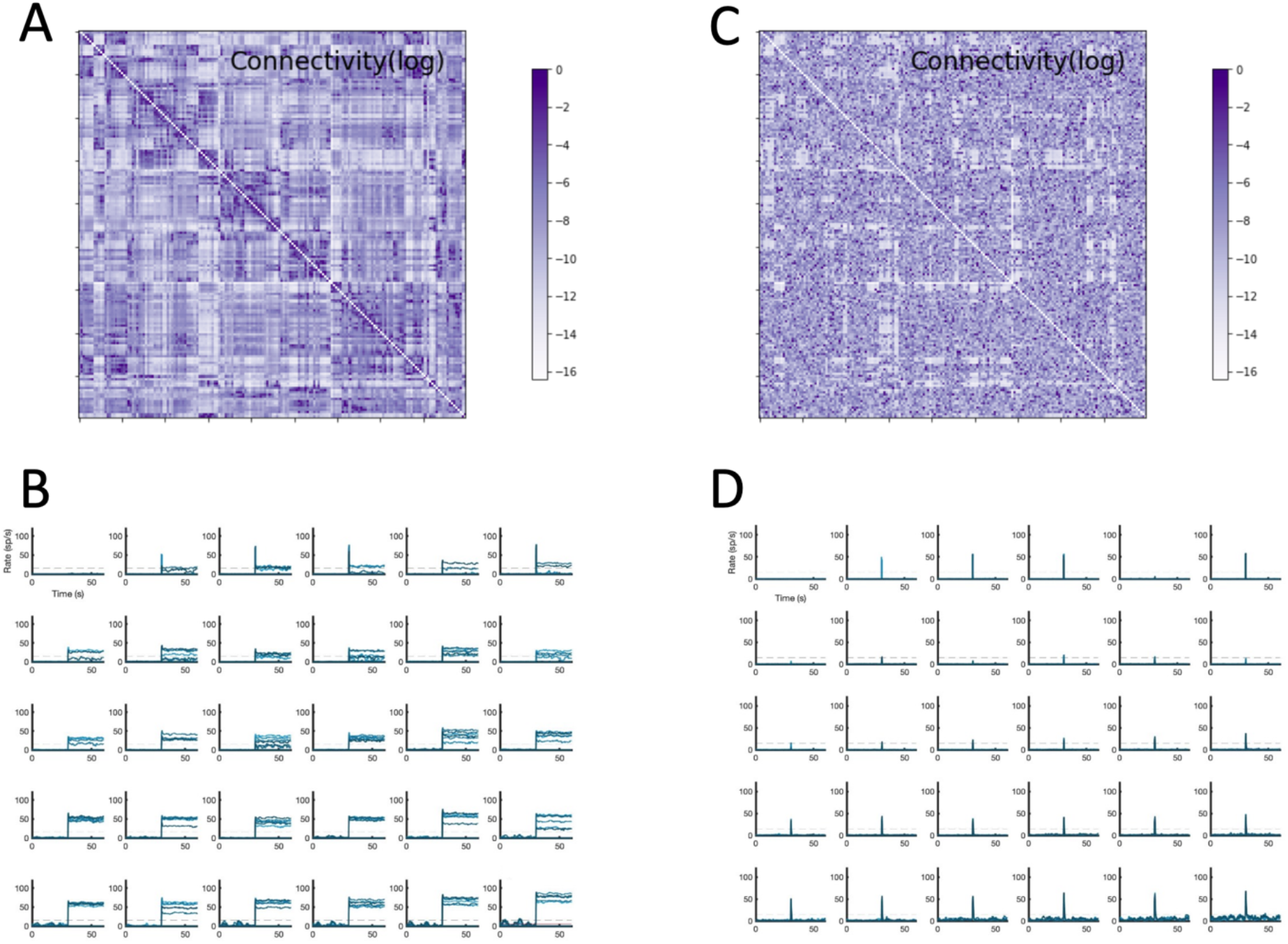
Distributed working memory in control and surrogate networks. (A) Connectivity matrix of the original model. (B) Firing rates as a function of time for all 180 areas in the model (each box displays six areas) in a working memory task as in Fig. 1 (C) Connectivity matrix for the surrogate network (weights are randomly swapped but the connectivity degree of each area is maintained). (D) Firing rates as a function of time for all areas in the surrogate network.

**Figure S3:**
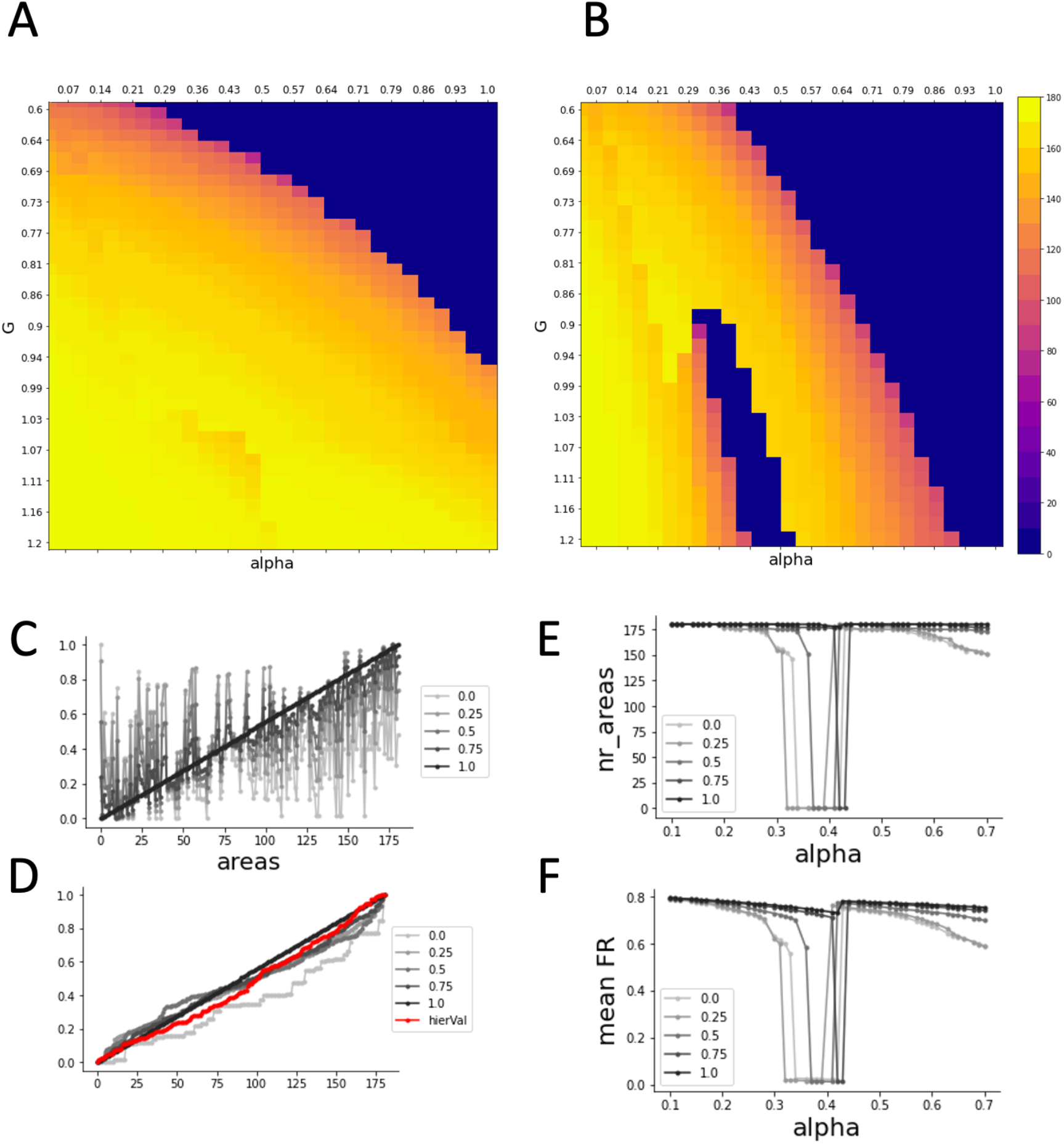
Effect of the heterogeneity induced by NMDAr density in the parameter space. (A) Activity map in the G-alpha parameter space, when T1w:T2w is used to constrain the local synaptic strength. (B) Activity map when NMDAr density is used to constrain local synaptic strength. (C) Linearisation of NMDA receptor density. The receptor densities are first sorted by hierarchy values. The value in the legend shows the strength of the linearization: the higher the value, the stronger is the linearization (completely linearised for 1) (D) NMDA with different linearization degree and T1w:T2w hierarchy sorted by values. This shows that T1w:T2w is much more linear and regular than NMDAr. (E) Number of areas with firing rates above 20 spikes/second for gradient with different linearity. (F) Normalized mean firing rates for the areas in panel E.

**Figure S4:**
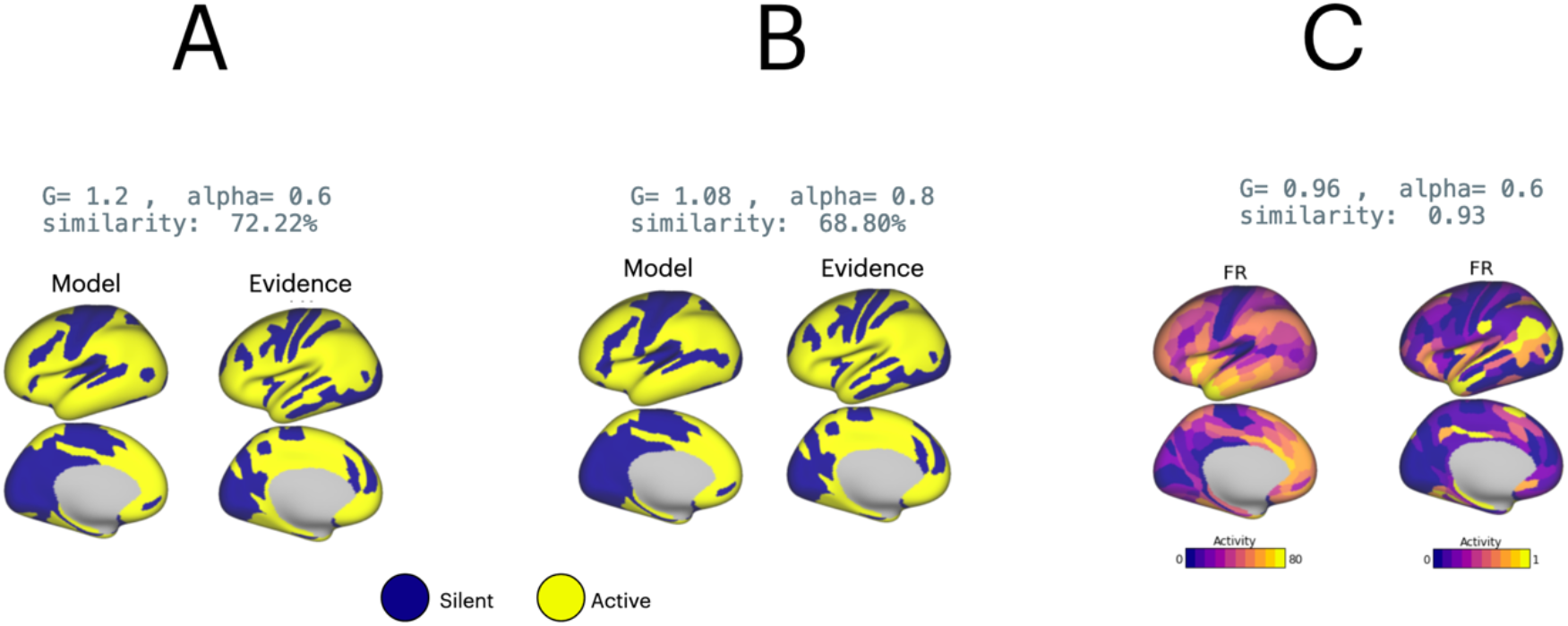
Similarity between evidence and model - three metrics. (A) Similarity measured by counting the number of overlapping areas. (B) Similarity measured by the overlapping surface areas. (C) Similarity measured by cross-correlation.

**Figure S5:**
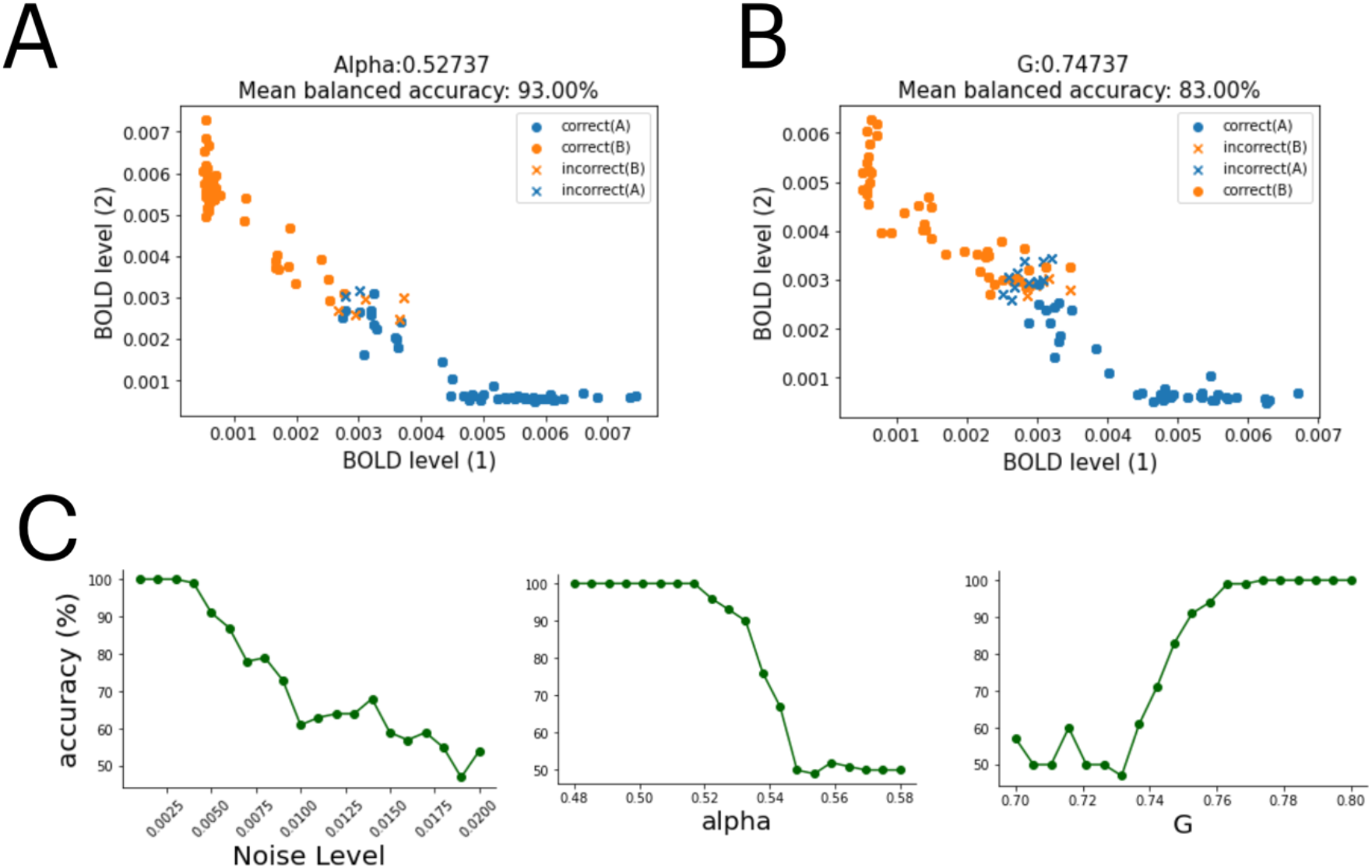
V1 Decoding of visual stimuli from mean BOLD signals for other conditions, and effects of parameters. (A) G = 0.75, alpha = 0.527, noise level = 0.01. (B) G = 0.747, alpha = 0.53, noise level = 0.01. Dots and crosses denote predictions of stimuli A (left) and stimuli B (right). (C) Effects of the noise level, feedback inhibition and global coupling on the decoding accuracy.

**Table S1:**
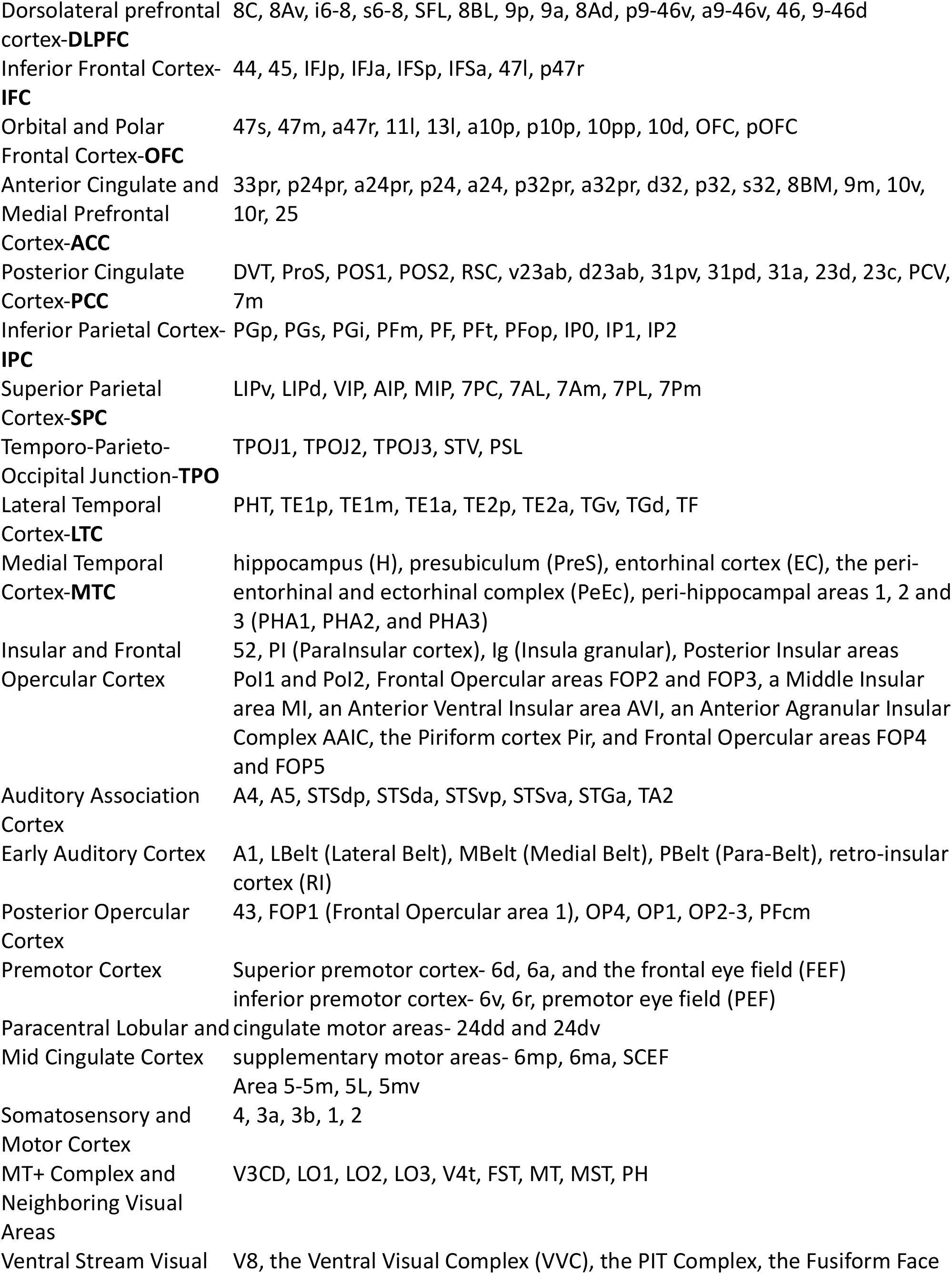
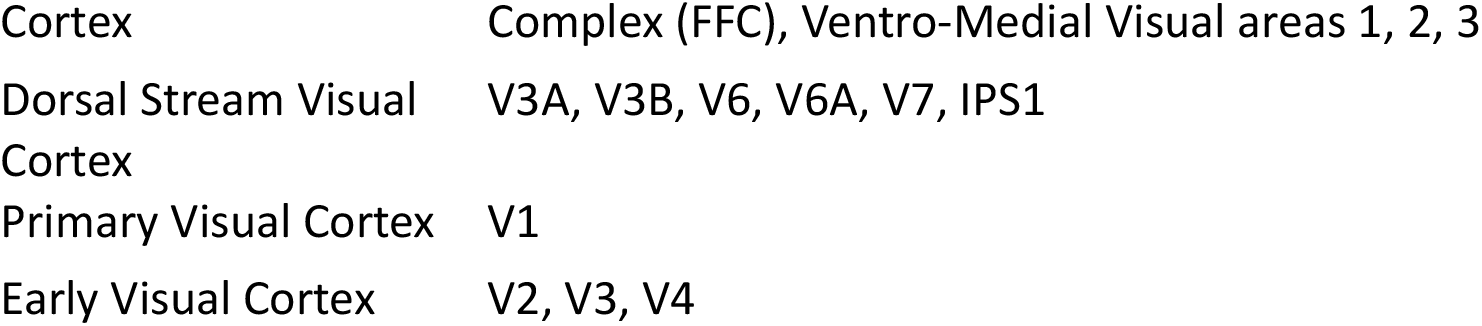
Parcellation adopted in this study

## Notes

### Competing Interest Statement

The authors have declared no competing interest.

